# The genetic relationship between female reproductive traits and six psychiatric disorders

**DOI:** 10.1101/433946

**Authors:** Guiyan Ni, Azmeraw Amare, Xuan Zhou, Natalie Mills, Jacob Gratten, Sang Hong Lee

## Abstract

Female reproductive behaviors have an important implication in evolutionary fitness and health of offspring. Previous studies have shown that age at first birth of women (AFB) is genetically associated with schizophrenia (SCZ). However, for most other psychiatric disorders and reproductive traits, the latent shared genetic architecture is largely unknown. Here we used the second wave of UK Biobank data (N=220,685) to evaluate the association between five female reproductive traits and polygenetic risk scores (PRS) projected from genome-wide association study summary statistics of six psychiatric disorders (N=429,178). We found that the PRS of attention-deficit/hyperactivity disorder (ADHD) were strongly associated with AFB (genetic correlation of −0.68 ± 0.03 with p-value = 1.86E-89), age at first sexual intercourse (AFS) (−0.56 ± 0.03 with p-value = 3.42E-60), number of live births (NLB) (0.36 ± 0.04 with p-value = 4.01E-17) and age at menopause (−0.27 ± 0.04 with p-value = 5.71E-13). There were also robustly significant associations between the PRS of eating disorder (ED) and AFB (genetic correlation of 0.35 ± 0.06), ED and AFS (0.19 ± 0.06), Major depressive disorder (MDD) and AFB (−0.27 ± 0.07), MDD and AFS (− 0.27 ± 0.03) and SCZ and AFS (−0.10 ± 0.03). Our findings reveal the shared genetic architecture between the five reproductive traits in women and six psychiatric disorders, which have a potential implication that helps to improve reproductive health in women, hence better child outcomes. Our findings may also explain, at least in part, an evolutionary hypothesis that causal mutations underlying psychiatric disorders have positive effects on reproductive success.

## INTRODUCTION

Female reproductive behaviours, including age at first birth (AFB), age at first sexual intercourse (AFS), age at menarche (AMC), age at menopause (AMP) and number of live births (NLB) have important implications in reproductive health and evolutionary fitness^1,2^. Some female reproductive traits have been associated with the physical and mental health of offspring^3^ and there has been growing evidence that maternal AFB is associated with increased risk of psychiatric disorder^4,5^ and behavioural problems in their children^6,7^. For instance, early or late maternal AFB is associated with offspring’s risk to schizophrenia (SCZ)^4^, bipolar disorder (BIP)^8^, attention-deficit/hyperactivity disorder (ADHD)^7^, autism spectrum disorder (ASD)^9^ and depression^10^ in offspring. Age at menopause and menarche also tend to be correlated with the risk of adverse mental health outcomes in offspring^11–14^.

The relationship between reproductive behavior and susceptibility to psychiatric disorders is likely complex and bidirectional. Individuals with psychiatric illness and their relatives may be more prone to risk taking and impulsive behavior, which cause early pregnancy and childbirth in women, or they may exhibit poor social interactions that result in delays in key reproductive transitions such as marriage, pregnancy and childbirth^15–20^.

In addition to the epidemiological evidence for a relationship between female reproductive traits and psychiatric disorder risk in offspring, studies have found a genetic basis for reproductive behavior traits^21,22^ and psychiatric disorders^23–25^. Genome wide association studies (GWAS) have identified several genome-wide significant genetic loci influencing reproductive behavior traits such as AFB^21^, NLB^21^.and AFS^3^. Similarly, GWASs on psychiatric disorders have successfully discovered a number of genome-wide significant genetic variants associated with the susceptibility of ADHD^26^, ASD^27^, eating disorders (ED)^28^, BIP^29^, major depressive disorder (MDD)^30^, and SCZ^31^. Therefore, it becomes important to estimate the shared genetic architecture between reproductive traits and psychiatric disorders. A few studies focused on understanding the complex genetic relationship between reproductive traits and psychiatric disorders^32^. For instance, recent studies have revealed genetic overlap between the risk of SCZ and AFB^23,25^. They showed a U-shaped relationship between AFB and the polygenic risk score of SCZ^23^, demonstrating that the epidemiological observation of a (phenotypic) association between maternal age and SCZ risk in offspring (e.g. McGrath et al.^4^, El-Saadi et al.^33^, Byrne et al. ^34^) could be partially explained by the latent genetic relationship^24^. However, for most other psychiatric disorders, the latent genetic architecture associated with reproductive traits is largely unknown. It would be interesting, for example, to estimate the extent of shared genetic architecture between reproductive traits and a wider range of psychiatric disorders. This would elucidate the latent relationship underlying the epidemiological observations of a phenotypic association between psychiatric disorders and reproductive traits^13,14^, which in turn contributes to improving reproductive health in women.

Here, we investigate the genetic association of five specific reproductive traits in women (AFB, AFS, AMC, AMP and NLB) with six common psychiatric disorders (ADHD, ASD, ED, BIP, MDD, and SCZ). This study will shed light on the complex bio-psychosocial risk factors associated with shared genetic effects between female reproductive behavior traits and a wide range of mental disorders. Furthermore, this will give important insight into how complex phenotypes of reproductive experience and mental health have been shaped through the interaction between multiple pleiotropic genes.

## MATERIALS AND METHODS

### Data

#### UK Biobank sample and quality control

We used the second wave of the UK Biobank dataset that initially included 264,859 women with imputed genotypes for 35.6 million SNPs. The imputation was based on the Haplotype Reference Consortium reference panel^35^. The quality control (QC) criteria for the imputed genotype data included an imputation reliability (INFO score) > 0.6, minor allele frequency > 0.01, p-value for Hardy-Weinberg equilibrium test > 10E-7, and SNP missingness < 0.05. Women with non-British ancestry or individual missingness > 0.05 or individuals with putative sex chromosome aneuploidy or one individual per pair of relatives was excluded. After the QC above, we compared discordance between the first and second wave of QCed UK Biobank data for each SNP and individual and excluded SNPs and individuals with a discordance rate above 0.05. In addition, we excluded genome-wide significant SNPs in a GWAS analysis where individuals in the first and second wave UK Biobank data were treated as cases and controls. Subsequently, 220,685 individuals and 7,253,311 SNPs remained, from which 1,133,064 Hapmap3 SNPs were selected for the analyses. Traits of interest were age at first birth (AFB), age at first sexual intercourse (AFS), age at menarche (AMC), age at menopause (AMP), and number of live births (NLB). Number of observations for each trait are provided in Table 1.

**Table 1.**
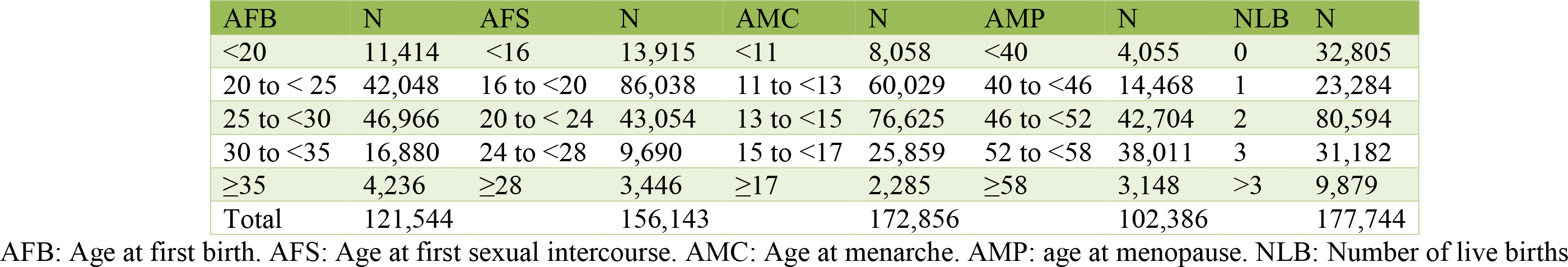
Sample breakdown by age at first birth, age first had sexual intercourse, age at menarche, age at menopause and number of live births

#### Psychiatric genomics consortium (PGC) GWAS summary statistics results

The GWAS summary results of six psychiatric disorders were obtained from the PGC (http://www.med.unc.edu/pgc), including attention-deficit/hyperactivity disorder (ADHD)^26^, autism spectrum disorder (ASD)^27^, eating disorder (ED)^28^, bipolar disorder (BIP)^29^, major depressive disorder (MDD)^30^, and schizophrenia (SCZ)^31^. The number of cases and controls in these studies are provided in Table S1.

### Statistical analyses

#### Estimation of polygenic risk scores (PRS)

For each psychiatric disorder, PRS were computed for the UK Biobank sample as the sum of the risk alleles weighted by the estimated SNP effects from the PGC GWAS that are in public domain (http://www.med.unc.edu/pgc/) and not likely to include UK Biobank sample. To compute PRS for the UK Biobank sample we used approximately1,133,064 HapMap 3 SNPs from each study. The projection of the SNP effects onto the UK Biobank data was conducted using MTG2^36^.

#### Mean difference of PRS across the five categories

We divided UK Biobank samples into five arbitrary categories according to their ages or the number of children (Table 1). For individuals in each category of each reproductive trait, we compared the mean difference for each of the six psychiatric disorders. P-values obtained based on two-tailed t-tests.

#### Linear prediction

Using a linear or polynomial regression model, we assessed, for each disorder, if PRS significantly predicted each of the phenotypes of the reproductive traits. In the prediction model, each of the reproductive traits was adjusted for age at interview, sex, year of birth, study center, genotype batch, and the first 15 principal components (PCs) provided by the UK Biobank. For sensitivity analyses, we repeated the prediction using additional variables including educational attainment, income level, and smoking and alcohol consumption status.

#### Genetic correlation

Genetic correlation is a classical population parameter to infer the geometric mean of trait variance (i.e. the additive genetic covariance between two traits scaled by the square root of the product of the genetic variance for each trait). Pairwise genetic correlations were estimated between the reproductive traits and psychiatric disorders using linkage disequilibrium score regression (LDSC) based on GWAS summary statistics^37,38^. GWAS was carried out and SNP effects were estimated for each of the reproductive traits using PLINK 1.9^39^. In the GWAS, phenotypes were adjusted for sex, age at interview, year of birth, assessment center at which participant consented, genotype batch and the first 15 PCs. We also used additional covariates such as educational attainment, income level, smoking and alcohol consumption status to further adjust the phenotypes in sensitivity analyses.

#### Analysis of overlapping samples between PGC and UK Biobank data

The intercept of LDSC is estimated as 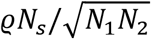, where ϱ is the phenotypic correlation among *N_S_* overlapping individuals in the two studies with sample sizes of *N*_1_ and *N*_2_, respectively^37^. We explicitly estimated the intercept when using LDSC to estimate genetic correlation between the reproductive traits and psychiatric disorders to check if there were any overlapping samples between UK Biobank and PGC.

## RESULTS

### Basic characteristics

In this study, we analyzed genetic data from 220,685 women who were part of the UK Biobank. The average age of the study participants at recruitment was 57, with a range from 40 to 71. The number of observations for each reproductive trait is provided in Table 1. The total sample of women was divided into five age categories according to their AFB, AFS, AMC, AMP or NLB status to detect the mean difference of PRS across the five categories. The age bin and the number of observations in each bin are provided in Table 1.

### Mean difference of PRS across the five categories

First, we computed PRS of the six psychiatric disorders for the UK Biobank sample and estimated the mean of the PRS for each age category of each reproductive trait (Figures 1-5). Then, the mean difference of the PRS was tested across the five categories for each of the reproductive traits (Tables S2-S6). A statistically significant difference in the PRS was determined after Bonferroni correction for multiple testing (i.e. significance level divided by the number of tests, 0.05/300=1.7E-4) and the results that passed the significance threshold were highlighted in bold (see Tables S2-S6).

**Figure 1.**
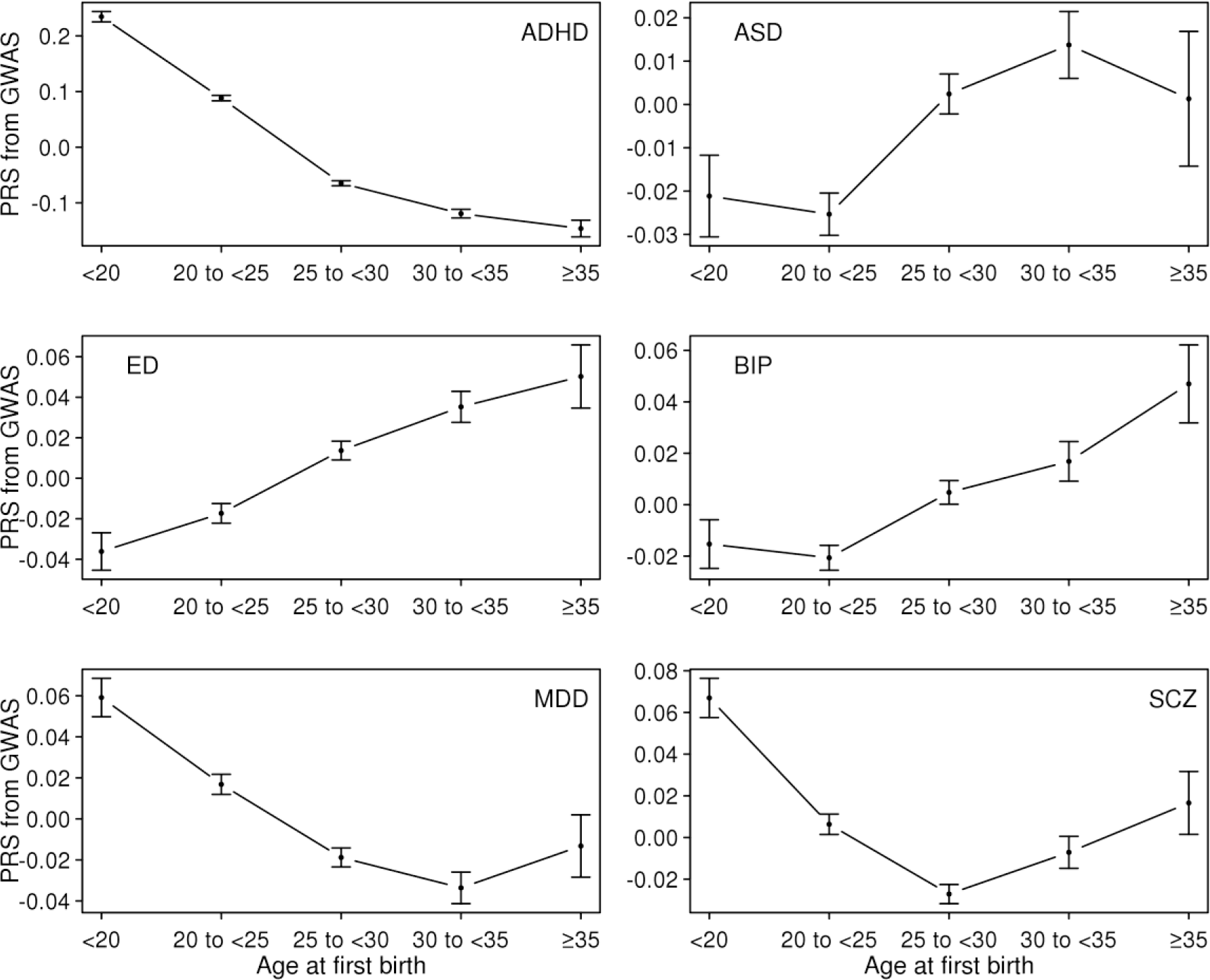
Means and standard errors of PRSs for the six psychiatric disorders, by age at first birth, in the UK Biobank sample.

In Figure 1, the direction of association between the PRS and AFB was positive (the older the AFB, the higher the risk) for ED, ASD and BIP, but negative (the younger the AFB, the higher the risk) for ADHD, MDD and SCZ although some traits (MDD and SCZ) showed non-linear (U-shaped) associations. A U-shaped relationship was previously shown between AFB and SCZ in an independent study^23^. The mean difference in PRS was statistically significant for most pairwise comparisons of age classes for ADHD, ED and MDD (Table S2). The significance was particularly pronounced for ADHD, with a P-value of 2.0E-184, for example, for the difference between women with AFB < 20 and women with 30 ≤ AFB <35 (Table S2). For SCZ, most significant signals came from younger AFB groups (Figure 1 and Table S2).

The results for AFS were similar to those for AFB, consistent with the strong correlation between these traits (Figure 6), although some signals were reduced and not significant after the Bonferroni correction (ASD, BIP and MDD) (Figure 2 and Table S3).

**Figure 2.**
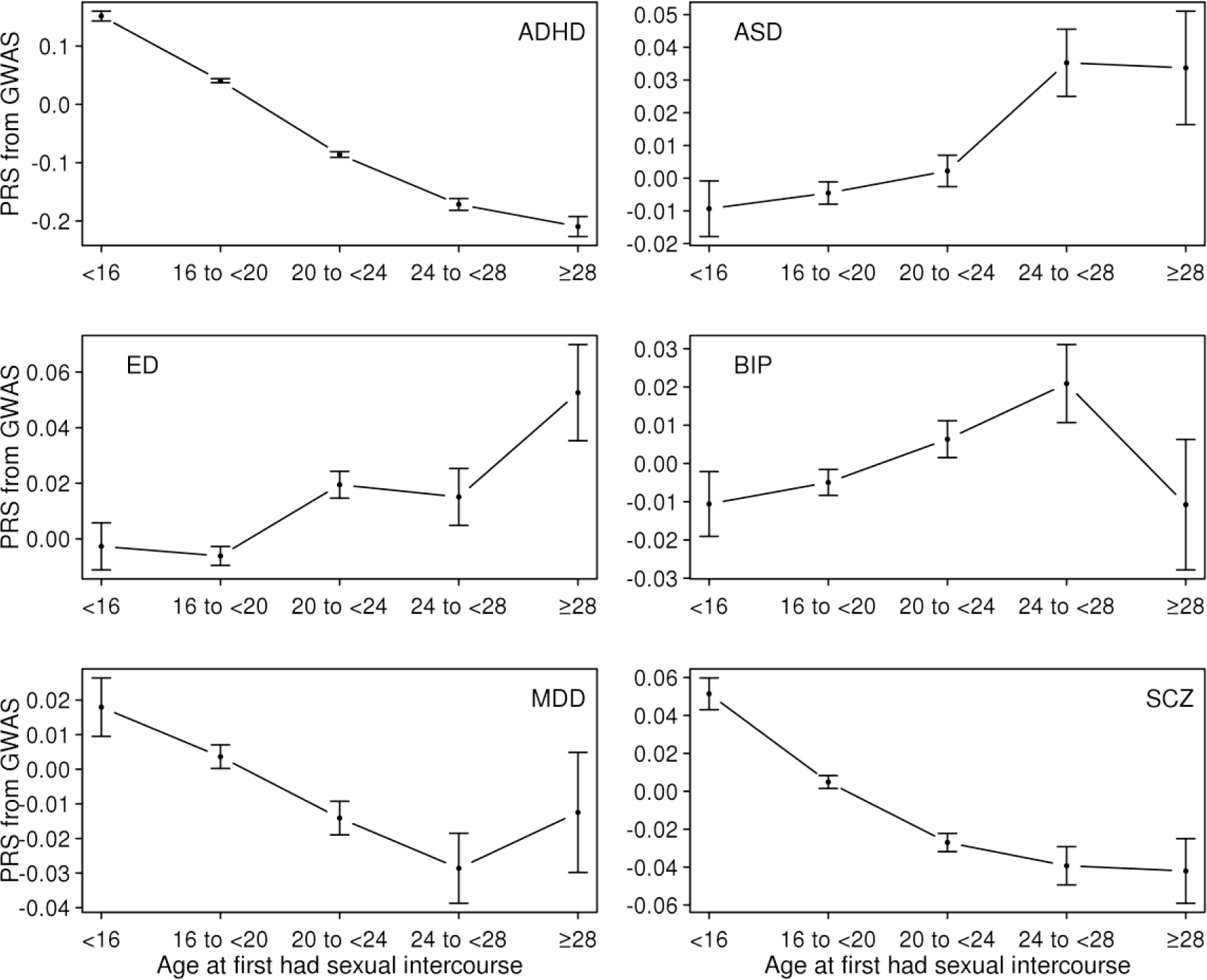
Means and standard errors of PRSs for the six psychiatric disorders, by age at first sexual intercourse, in the UK Biobank sample.

For AMC and AMP, there were relatively few statistically significant differences between age groups for the six psychiatric disorders, with the exception of ADHD (Tables S4 and S5). The relationship between the PRS of ADHD and AMC was non-linear (Figure 3), with earlier or later AMC tending to have significantly higher ADHD risk than intermediate AMC (Table S4). A similar relationship was observed between the ADHD PRS and AMP in that earlier or later AMP was associated with higher ADHD risk than moderate AMP (Figure 4 and Table S5).

**Figure 3.**
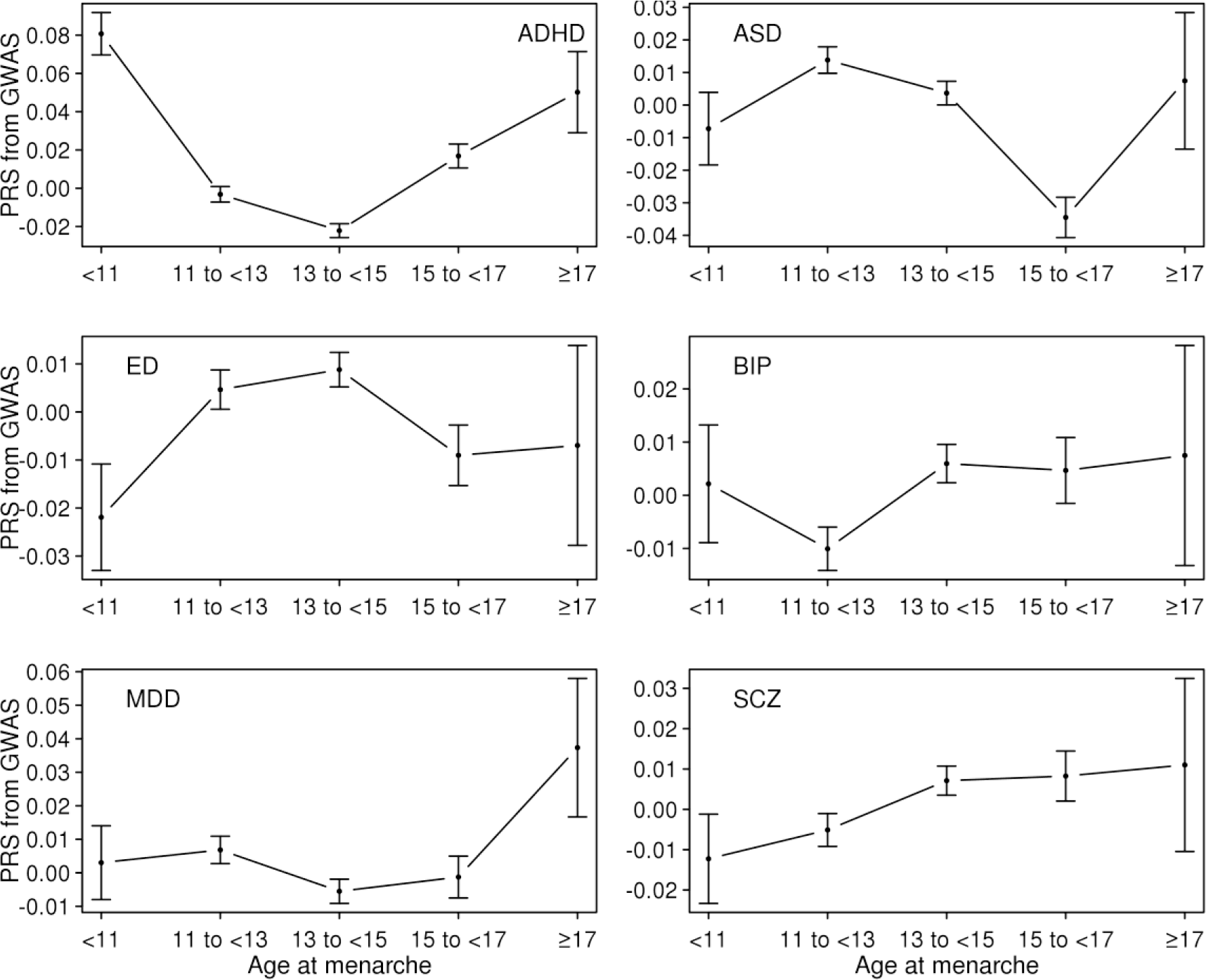
Means and standard errors of PRSs for the six psychiatric disorders by age at menarche in the UK Biobank sample.

**Figure 4.**
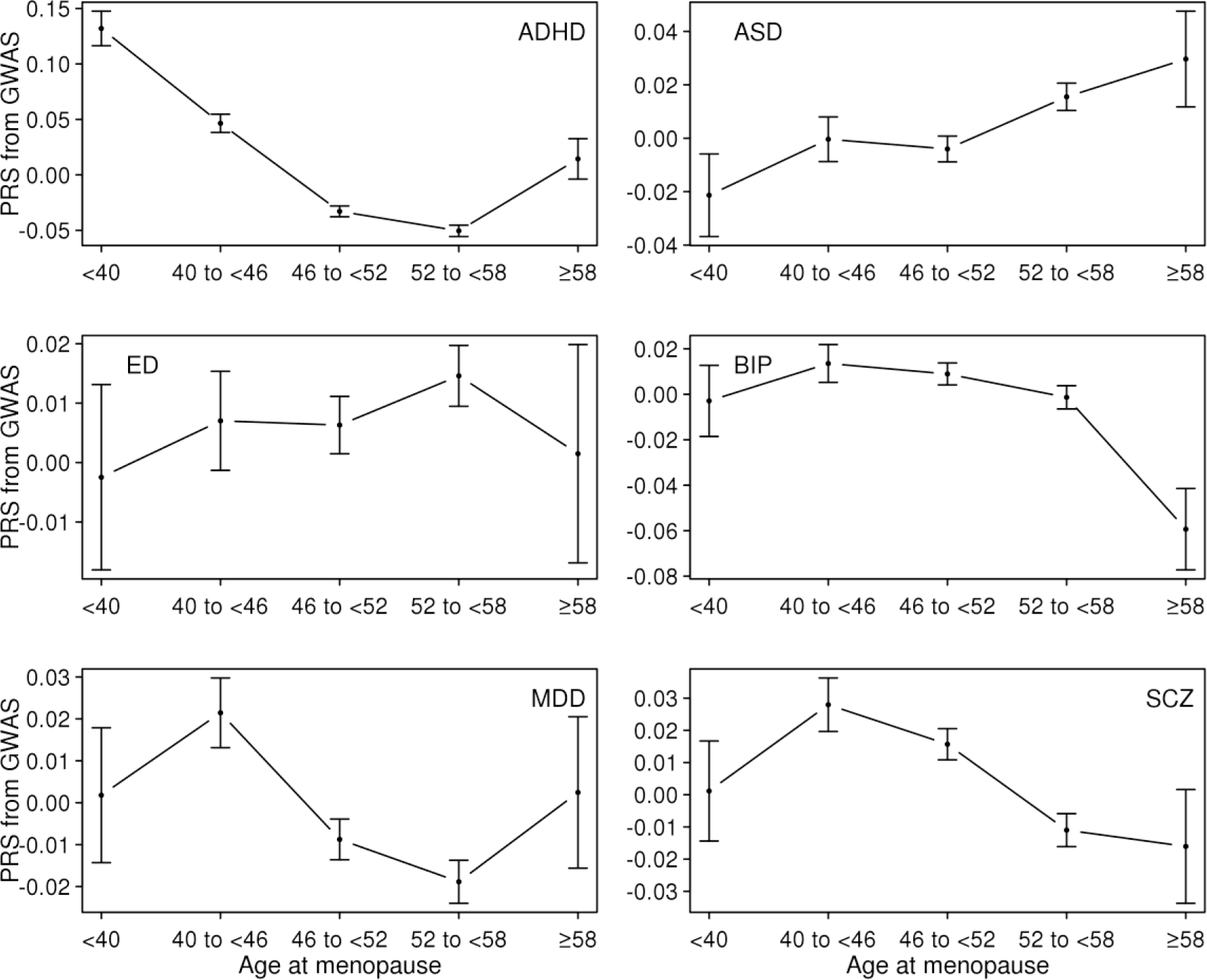
Means and standard errors of PRSs for the six psychiatric disorders by age at menopause in the UK Biobank sample.

Since NLB was expected to be negatively correlated with AFB (see Figure 6), the pattern of the mean PRS of ADHD was inversely correlated between the groups classified according to NLB and AFB (Figures 1 and 5). Table S6 shows that the ADHD PRS was strongly associated with NLB. In addition, there were a number of significant association signals of NLB for MDD and SCZ (Table S6).

**Figure 5.**
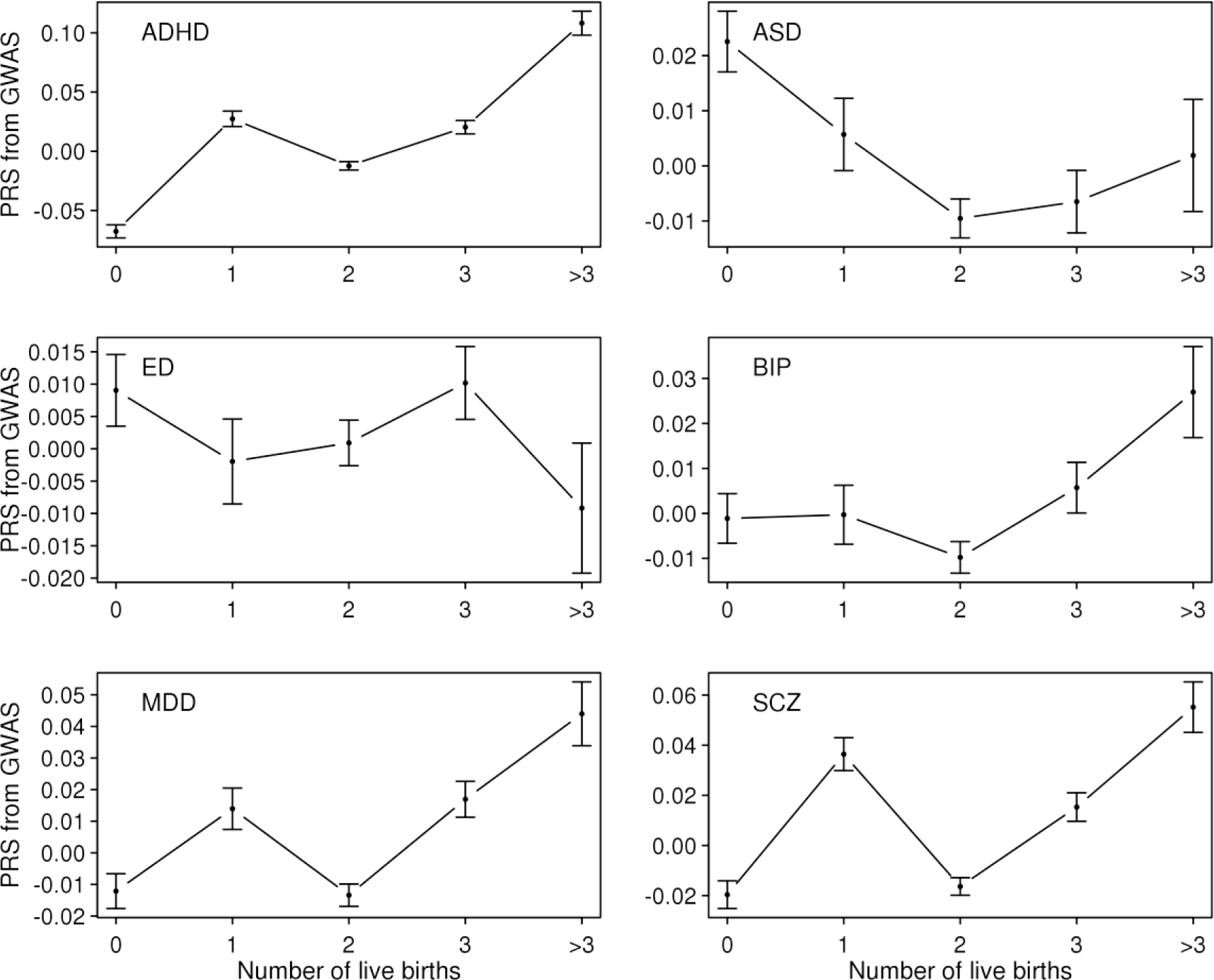
Means and standard errors of PRSs for the six psychiatric disorders by number of live births in the UK Biobank sample.

**Figure 6.**
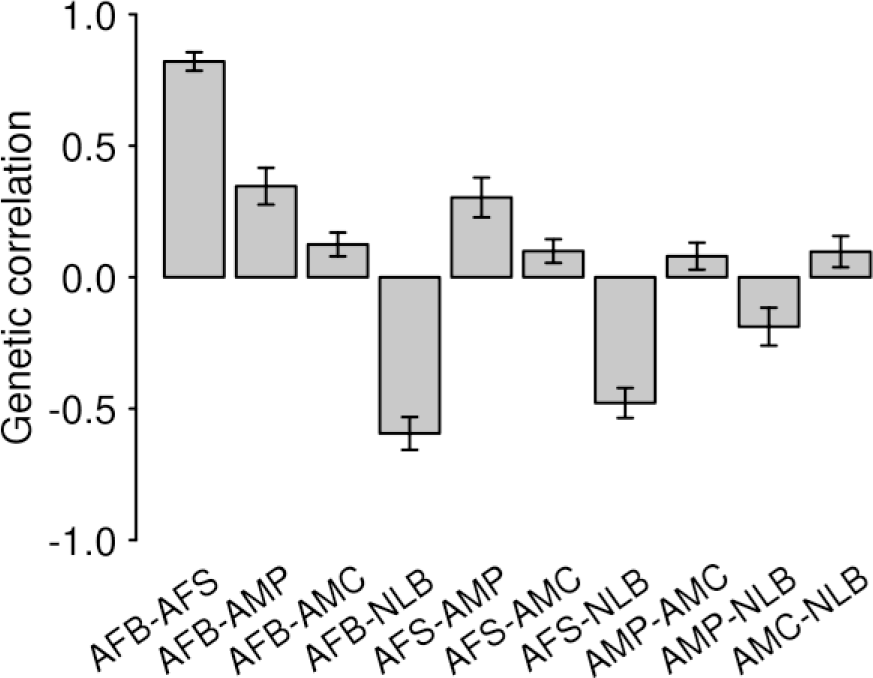
Genetic correlations among the five reproductive traits estimated using the base model. In the base model, the reproductive traits were adjusted for age at interview, sex, year of birth, study center, genotype batch, and the first 15 principal components. Error Bars are 95% confidence intervals. AFB: Age at first birth. AFS: Age at first sexual intercourse. AMC: Age at menarche. AMP: age at menopause. NLB: Number of live births

### Linear prediction

We used a regression method to assess the significance of genetic association between the reproductive behavior traits and six psychiatric disorders (see Methods).

We regressed each of pre-adjusted reproductive traits (AFB, AFS, AMC, AMP and NLB) on the PRS of each psychiatric disorder. Using the linear regression prediction modeling, we showed that the PRS of each six psychiatric disorder were significantly associated with at least one of the five reproductive traits, confirming some of the robust associations from the earlier analyses of mean difference of PRS across the five age categories (Tables S2-S6). Of all the five reproductive traits, AFB was the trait best predicted by the PRS of the six psychiatric disorders (Figure 7, the corresponding R-squared and P-values are in Table S7). Because AFB and AFS are highly correlated traits, the results for AFS were similar to those for AFB except that there was no significant association between PRS of AFS and BIP, which is consistent with the analyses of mean difference of PRS above (Tables S2 and S3). NLB was significantly predicted by the PRS of ADHD, ASD, MDD, and SCZ. In addition, AMC and AMP were only associated with the PRS of ASD and ADHD, respectively. We noted that the majority of significance was explained by linear predictions, but not by quadratic polynomial predictions for all of the associations (Table S8). There were marginal significances for quadratic polynomial associations only for a few pairwise comparisons (AFB vs SCZ and AFS vs ED).

**Figure 7.**
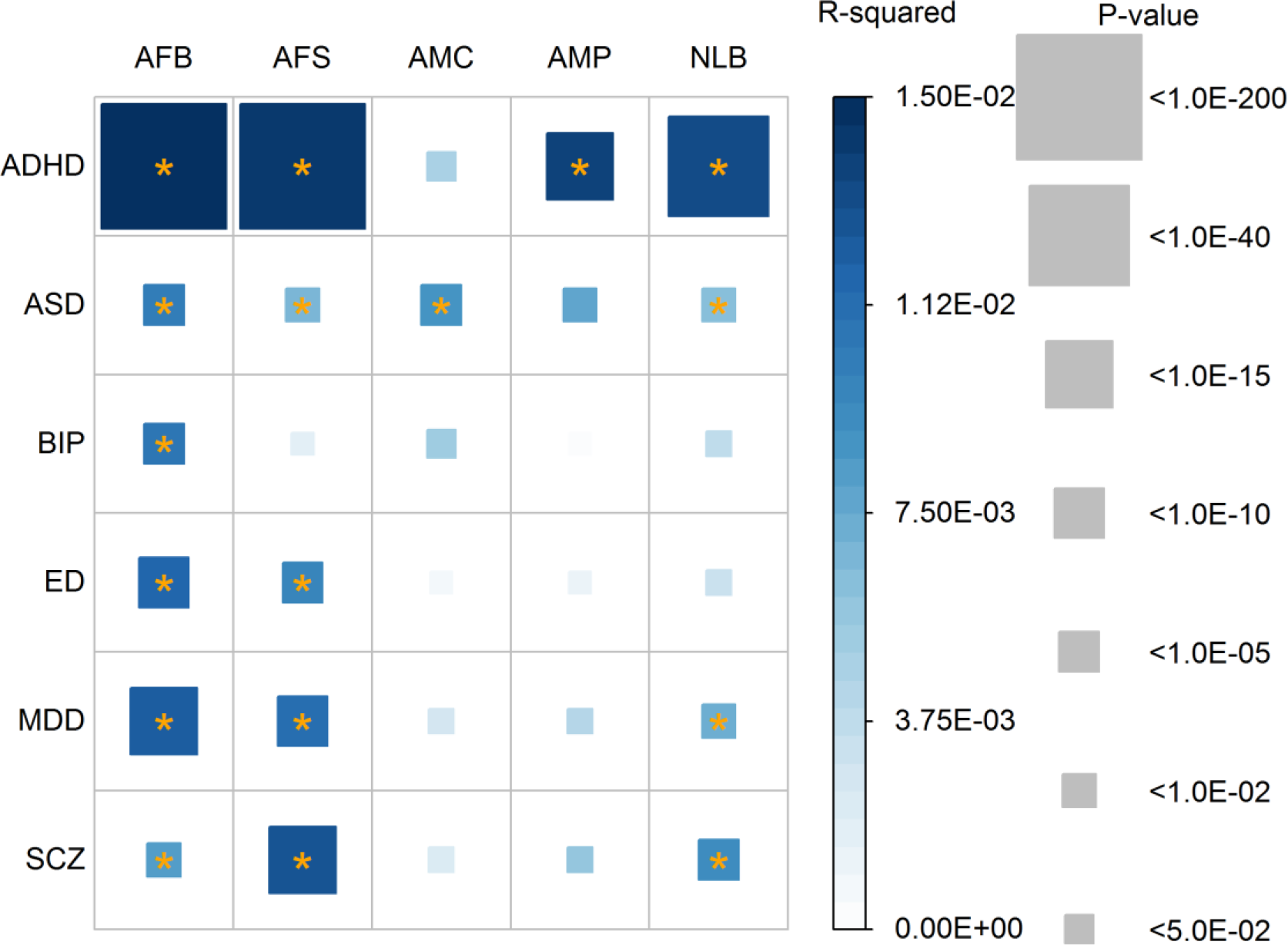
Coefficient of determination (R^2^) and p-values for its significance based on a linear prediction model. Color of each box represents the level of R-squared, and the size of squares represents its significance (p-value). R-squared that are significantly different from zero after Bonferroni correction (0.05/30) are marked with an asterisk. Dependent variables were adjusted for age at interview, year of birth, assessment centre at which the participant consented, genotype batch, and the first 15 principal components. The number of records used for the analyses was 121,544 for AFB, 156,143 for AFS, 102,386 for AMP, and 172,856 for AMC and 177,744 for NLB. AFB: age at first firth. AFS: age at first sexual intercourse. AMP: age at menopause. AMC: age at menarche. NLB: number of live births. ADHD: Attention-Deficit/Hyperactivity Disorder. ASD: Autism spectrum disorder. ED: Eating disorder. BIP: Bipolar disorder. MDD: Major depressive disorder. SCZ: Schizophrenia

We conducted a sensitivity analysis in which dependent variables were further adjusted for educational attainment, income level, smoking and alcohol consumption status (Figure S1 and Table S9). Most of the association signals remained significant with some exceptions. For example, the association of AFB, AFS, and NLB with ASD disappeared, as did the association between AFB and BIP. Conversely, ADHD became significantly associated with AMC after correcting for the additional covariates.

### Genetic correlations

We used LDSC to estimate genetic correlations between the reproductive behavior traits and six psychiatric disorders (see Methods). We estimated genetic correlations between the five reproductive traits to reveal the shared genetic architecture of the traits (Figure 6). As expected, the genetic correlation between AFB and AFS was very high (0.821± 0.018) and that between AFB and NLB was high (−0.594 ± 0.032). However, the genetic correlation between AMC and AMP was relatively low and they also have moderate or low genetic correlations with other reproductive traits, e.g. 0.346 ± 0.036 between AFB and AMP, 0.303 ± 0.039 between AFS and AMP and ~ 0.1 between AMC and other traits (Figure 6). These results explain the observations in the analyses of mean difference in PRS (Figures 1–5) and linear predictors (Figure 7), where the results between AFB and AFS were mostly similar, and those between AFB (AFS) and NLB were reciprocally similar for the association with ADHD PRS. The estimated genetic correlations among those five reproductive traits remained similar after additional adjustment of the dependent variables for educational attainment, income levels, and smoking and alcohol consumption status (Figure 6 vs. Figure S2).

Figure 8 shows the estimated genetic correlation from LDSC for each pair of five reproductive traits and six psychiatric disorders. The detailed genetic correlations and P-value are in Table S10. Nine pairs of genetic correlations out of 30 were significantly different from zero after Bonferroni correction. For AFB analyses, the estimated genetic correlations were greater than zero (positive association) between AFB and ED (0.349 ± 0.061), and lower than zero (negative association) between AFB and ADHD (−0.677 ± 0.034) and AFB and MDD (−0.273 ± 0.069). Similarly, AFS was inversely correlated with ADHD (−0.563 ± 0.034), MDD (−0.265 ± 0.066) and SCZ (−0.100 ± 0.030) and positively correlated with ED (0.189 ± 0.055). For AMC and AMP, there was no significant genetic correlation except that between AMP and ADHD (−0.272 ± 0.038). NLB showed positive genetic correlation with ADHD (0.356 ± 0.042) and was non-significant for other pairs of traits. These results agreed with the analyses of mean difference of PRS (Figures 1-5) and linear prediction above (Figure 7).

**Figure 8.**
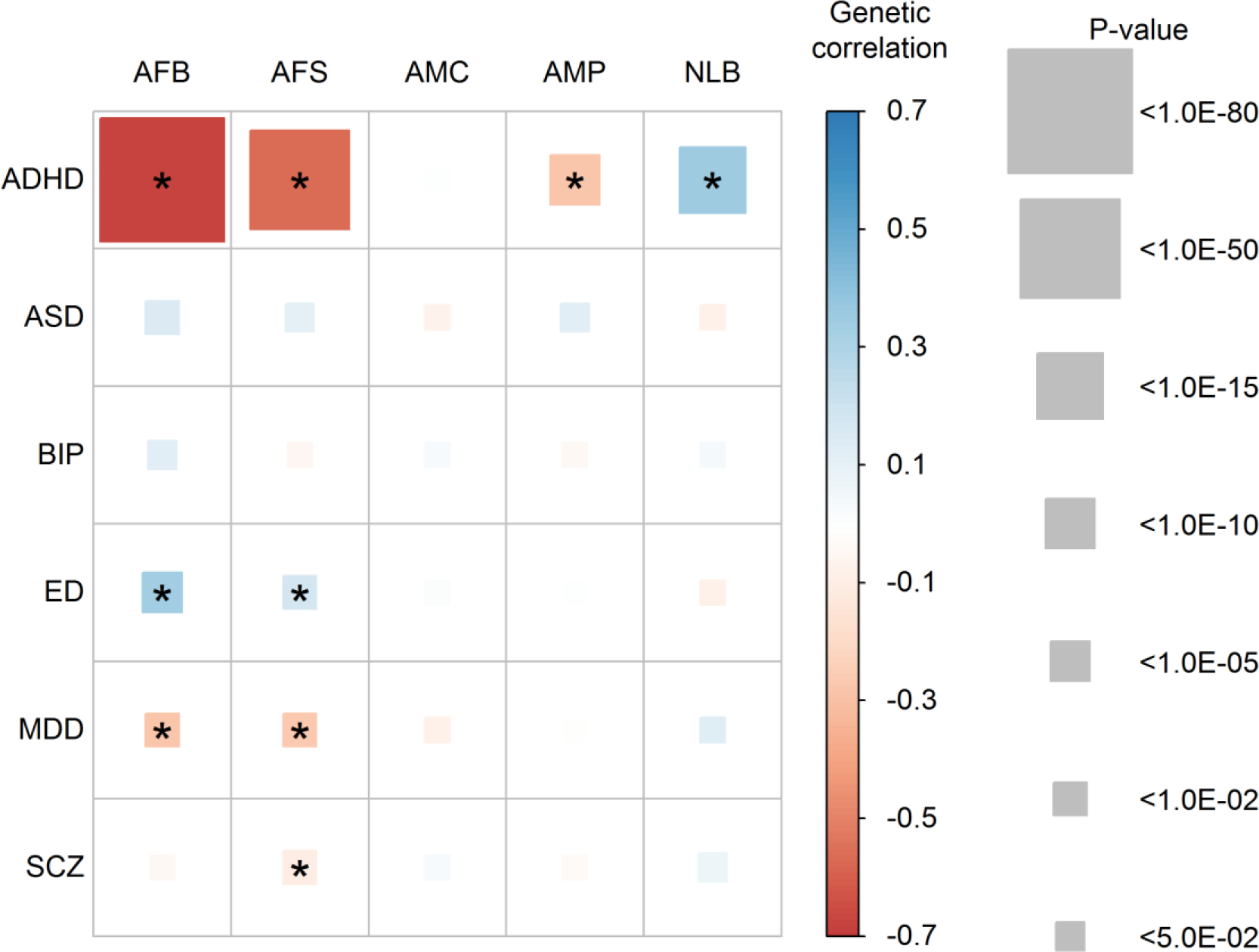
Genetic correlations between the five reproductive traits and the six psychiatric disorders estimated using the base model. Color of each box represents the level of estimated genetic correlation (blue for positive and red for negative correlation), and the size of squares represents its significance (p-value). Estimated genetic correlations that are significantly different from zero after Bonferroni correction (0.05/30) are marked with an asterisk. AFB: Age at first birth. AFS: Age at first sexual intercourse. AMC: Age at menarche. AMP: age at menopause. NLB: Number of live births ADHD: Attention-Deficit/Hyperactivity Disorder. ASD: Autism spectrum disorder. ED: Eating disorder. BIP: Bipolar disorder. MDD: Major depressive disorder. SCZ: Schizophrenia In the base model, the reproductive traits were adjusted for age at interview, sex, year of birth, study center, genotype batch, and the first 15 principal components.

In the analyses where dependent variables were further adjusted for education, income levels, smoking and alcohol consumption status, the estimated genetic correlations between reproductive traits and psychiatric disorders were not substantially changed (Figure S3 and Table S10), compared to those depicted in Figure 8 and Table S10.

### Analysis of overlapping samples between PGC and UK Biobank data

The intercepts between the reproductive traits and six psychiatric disorders were not significantly different from zero, indicating that there were no overlapping samples between UK Biobank and PGC (Figures S4 and S5). The sole exception was that the intercept between AFB and ED was significantly lower than zero (−0.017 ± SE 0.006), which may be due to sampling errors or excessive heterogeneity^37,38^. Most of the intercepts from the cross-trait LDSC analyses of the five reproductive traits were significantly different from zero, as expected (Figures S6 and S7). We note that the intercepts of AMP-AMC and AMC-NLB were not different from zero, which was because of the small phenotypic correlation between these pairs of traits.

## DISCUSSION

We revealed the complex psychosocial genetic risk architecture underpinning the association between reproductive traits and six psychiatric disorders. The strong genetic associations between ADHD and most reproductive traits (except AMC) were supported by three different approaches, the analyses of mean difference of PRS across five different (age) categories, linear prediction and genetic correlation using LDSC. Genetic associations between two of the reproductive traits (AFB and AFS) and both MDD and ED were consistently significant across the various analyses. The association between SCZ and AFB (and NLB) was significant in the analyses of mean difference of PRS and linear prediction although the estimated genetic correlation from LDSC analysis was not significant.

It is known that modelling genetic correlation between traits can increase accuracy significantly in predicting individual genetic risk for the traits^40,41^. Our findings of significant associations between female reproductive traits and between these traits and psychiatric disorders may be useful in improving female reproductive health, hence better child outcomes. For example, the high-risk group for ADHD could be informed about features of ADHD, such as impulsive behaviour and possible consequences of impulsivity. This intervention may lead to prevent them giving birth at an immature age, which can improve their reproductive health and maternal environment for their baby. Furthermore, this information of genetic predisposition, combined with information on psychosocial factors, can be recorded as a form of family medical history, and used to monitor health of offspring.

Psychiatric disorders are highly heritable traits (e.g. ADHD or SCZ) while they persist in the population in a stable prevalence rate. It is questioned why natural selection has not excluded the causal mutations underlying psychiatric disorders. There are some plausible hypotheses. First, the impact of natural selection on the removal of existing causal mutations may be slower than the addition of new genetic mutations causing, for example, ADHD or SCZ. Second, pre-existing neutral mutations may interact with environments that have been newly exposure to the population, and cause the disorders^42^. Third, causal mutations underlying psychiatric disorders have positive effects on reproductive success. The last hypothesis can be supported by our observation that NLB and ADHD have a strong positive genetic correlation (Figures 7 and 8). For SCZ and MDD, there is a suggestive signal for a positive genetic association with NLB (Figures 7 and 8). However, the hypothesis would not be supported by the results for ASD, BIP and ED.

We report a novel shared genetic architecture between female reproductive traits, i.e. including strong positive genetic correlation between AFB and AFS, high negative genetic correlation between AFB and NLB, and AFS and NLB, moderate positive genetic correlation between AMP and AFB (AFS), moderate negative genetic correlation between AMP and NLB, and negligible genetic correlation between AMC and other reproductive traits. These reported relationships among the reproductive traits may help to shed light on how causal genetic variants are involved and shared in the regulation of reproductive mechanisms at the genome-wide level. The strong positive and negative associations between AFB and AFS, and AFB and NLB are intuitive and well expected. However, other relationships (e.g. moderate positive association between AFB and AMP and negligible association between AMC and AMP) should be further investigated in a future study to identify the exact mechanism of menarche and menopause in relation to reproductive success^43^.

There are a number of limitations in this study. Firstly, we used the PRS or LDSC approach using GWAS summary statistics without having access to individual-level genotype data for psychiatric disorders, for which we do not have permissions. The power and accuracy of detection and estimation in the analyses could be increased when using an approach based on individual-level genotype data^44^. Secondly, we tested and estimated genetic association between the PRSs of six psychiatric disorders and five reproductive traits in women, which was underlain by pleiotropic effects. However, we did not determine if genetic variation for psychiatric disorders is causally related to these reproduction traits or vice versa. Thirdly, we only focused on reproductive traits in women because data on related male traits (e.g. AFB, AFS) are not fully available in the UKB. The collection of data on reproductive traits such as AFB in men, that would enable the study of genetic associations between psychiatric disorders and these reproductive traits should be a priority for the field.

In conclusion, we revealed the latent genetic architecture between the five reproductive traits in women and six psychiatric disorders. In particular, ADHD and a set of reproductive traits were substantially associated through common genetic factors. There were also a number of robust associations between reproductive traits and ED, SCZ and MDD. Our findings can have potential to help improve reproductive health in women and their child outcomes. These findings also can help address an evolutionary hypothesis that causal mutations underlying ADHD, MDD and SCZ have positive effects on reproductive success.

## URLS

PGC: http://www.med.unc.edu/pgc/

MTG2: https://sites.google.com/site/honglee0707/mtg2

LDSC: https://github.com/bulik/ldsc

## AUTHOR CONTRIBUTION

S.H.L. conceived the idea and directed the study. G.N. and S.H.L. performed the analyses and interpreted the results. J.G. and N.M. provided critical feedback and key elements in interpreting the results. S.H.L., G.N., J.G. and X.Z. drafted the manuscript. With support from S.H.L., A.A. made interpretation of the analyses output, organised the scientific theory of the work and drafted the manuscript. All authors contributed to editing and approval of the final manuscript.

## ACKNOWLEDGEMENTS

This research is supported by the Australian National Health and Medical Research Council (1080157, 1087889), and the Australian Research Council (DP160102126, FT160100229). This research has been conducted using the UK Biobank Resource. UK Biobank (http://www.ukbiobank.ac.uk) Research Ethics Committee (REC) approval number is 11/NW/0382. Our reference number approved by UK Biobank is 14575. This research was undertaken with the assistance of resources and services from the National Computational Infrastructure (NCI), which is supported by the Australian Government.

## REFERENCES

1. Tropf, F.C. et al. Hidden heritability due to heterogeneity across seven populations. Nature human behaviour 1, 757 (2017).

2. Tropf, F.C. et al. Human fertility, molecular genetics, and natural selection in modern societies. PLoS ONE 10, 1–14 (2015).

3. Day, F.R. et al. Physical and neurobehavioral determinants of reproductive onset and success. Nature genetics 48, 617 (2016).

4. McGrath, J.J. et al. A comprehensive assessment of parental age and psychiatric disorders. JAMA psychiatry 71, 301–9 (2014).

5. Croen, L.A., Najjar, D.V., Fireman, B. & Grether, J.K. Maternal and paternal age and risk of autism spectrum disorders. Archives of pediatrics & adolescent medicine 161, 334–340 (2007).

6. Barclay, K. & Myrskylä, M. Maternal age and offspring health and health behaviours in late adolescence in Sweden. SSM-population health 2, 68–76 (2016).

7. Chang, Z. et al. Maternal age at childbirth and risk for ADHD in offspring: A population-based cohort study. International Journal of Epidemiology 43, 1815–1824 (2014).

8. Menezes, P.R. et al. Paternal and maternal ages at conception and risk of bipolar affective disorder in their offspring. Psychological medicine 40, 477–485 (2010).

9. Idring, S. et al. Parental age and the risk of autism spectrum disorders: findings from a Swedish population-based cohort. International journal of epidemiology 43, 107–115 (2014).

10. Tearne, J.E. et al. Older maternal age is associated with depression, anxiety, and stress symptoms in young adult female offspring. Journal of abnormal psychology 125, 1 (2016).

11. Leibenluft, E. Women with bipolar illness: clinical and research issues. The American journal of psychiatry 153, 163 (1996).

12. Ballinger, C.B. Psychiatric aspects of the menopause. The British Journal of Psychiatry 156, 773–787 (1990).

13. Tondo, L., Pinna, M., Serra, G., De Chiara, L. & Baldessarini, R.J. Age at menarche predicts age at onset of major affective and anxiety disorders. European Psychiatry 39, 80–85 (2017).

14. Platt, J.M., Colich, N.L., McLaughlin, K.A., Gary, D. & Keyes, K.M. Transdiagnostic psychiatric disorder risk associated with early age of menarche: A latent modeling approach. Comprehensive Psychiatry (2017).

15. Özcan, N.K., Boyacıoğlu, N.E., Enginkaya, S., Dinç, H. & Bilgin, H. Reproductive health in women with serious mental illnesses. Journal of clinical nursing 23, 1283–1291 (2014).

16. Matevosyan, N.R. Reproductive health in women with serious mental illnesses: a review. Sexuality and Disability 27, 109–118 (2009).

17. Bellack, A.S., Morrison, R.L., Wixted, J.T. & Mueser, K.T. An analysis of social competence in schizophrenia. The British Journal of Psychiatry 156, 809–818 (1990).

18. Giordano, G.N., Ohlsson, H., Sundquist, K., Sundquist, J. & Kendler, K.S. The association between cannabis abuse and subsequent schizophrenia: a Swedish national co-relative control study. Psychological medicine 45, 407–414 (2015).

19. Salas-Wright, C.P., Vaughn, M.G., Ugalde, J. & Todic, J. Substance use and teen pregnancy in the United States: evidence from the NSDUH 2002– 2012. Addictive behaviors 45, 218–225 (2015).

20. Reddy, L.F. et al. Impulsivity and risk taking in bipolar disorder and schizophrenia. Neuropsychopharmacology 39, 456 (2014).

21. Barban, N. et al. Genome-wide analysis identifies 12 loci influencing human reproductive behavior. Nature genetics 48, 1462 (2016).

22. Perry, J.R.B. et al. Parent-of-origin-specific allelic associations among 106 genomic loci for age at menarche. Nature 514, 92 (2014).

23. Mehta, D. et al. Evidence for genetic overlap between schizophrenia and age at first birth in women. JAMA Psychiatry 73, 497–505 (2016).

24. Gratten, J. et al. Risk of psychiatric illness from advanced paternal age is not predominantly from de novo mutations. Nature Genetics 48, 718–724 (2016).

25. Ni, G., Gratten, J., Wray, N.R. & Lee, S.H. Age at first birth in women is genetically associated with increased risk of schizophrenia. Scientific reports 8, 10168 (2018).

26. Demontis, D. et al. Discovery Of The First Genome-Wide Significant Risk Loci For ADHD. bioRxiv (2017).

27. The Autism Spectrum Disorders Working Group of The Psychiatric Genomics, C. Meta-analysis of GWAS of over 16,000 individuals with autism spectrum disorder highlights a novel locus at 10q24.32 and a significant overlap with schizophrenia. Molecular Autism 8, 21 (2017).

28. Duncan, L. et al. Significant locus and metabolic genetic correlations revealed in genome-wide association study of anorexia nervosa. American Journal of Psychiatry 174, 850–858 (2017).

29. Psychiatric Gwas Consortium Bipolar Disorder Working Group. Large-scale genome-wide association analysis of bipolar disorder identifies a new susceptibility locus near ODZ4. Nature genetics 43, 977–983 (2011).

30. Ripke, S. et al. A mega-analysis of genome-wide association studies for major depressive disorder. Molecular psychiatry 18, 497–511 (2013).

31. Schizophrenia Working Group of the Psychiatric Genomics Consortium. Biological insights from 108 schizophrenia-associated genetic loci. Nature 511, 421–427 (2014).

32. Mullins, N. et al. Reproductive fitness and genetic risk of psychiatric disorders in the general population. Nature communications 8, 15833 (2017).

33. El-Saadi, O. et al. Paternal and maternal age as risk factors for psychosis: findings from Denmark, Sweden and Australia. Schizophrenia Research 67, 227–236 (2004).

34. Byrne, M., Agerbo, E., Ewald, H., Eaton, W.W. & Mortensen, P.B. Parental age and risk of schizophrenia: a case-control study. Archives of general psychiatry 60, 673–678 (2003).

35. Loh, P.-R. et al. Reference-based phasing using the Haplotype Reference Consortium panel. Nature genetics 48, 1443 (2016).

36. Lee, S.H. & van der Werf, J. MTG2: An efficient algorithm for multivariate linear mixed model analysis based on genomic information. Bioinformatics 32, 1420–1422 (2016).

37. Bulik-Sullivan, B. et al. An Atlas of Genetic Correlations across Human Diseases and Traits. Natrure genetics 47, 1236–1241 (2015).

38. Bulik-Sullivan, B.K. et al. LD Score regression distinguishes confounding from polygenicity in genome-wide association studies. 47, 291–295 (2015).

39. Chang, C.C. et al. Second-generation PLINK: rising to the challenge of larger and richer datasets. GigaScience 4(2015).

40. Maier, R. et al. Joint Analysis of Psychiatric Disorders Increases Accuracy of Risk Prediction for Schizophrenia, Bipolar Disorder, and Major Depressive Disorder. The American Journal of Human Genetics 96, 283–294 (2015).

41. Maier, R.M. et al. Improving genetic prediction by leveraging genetic correlations among human diseases and traits. Nature communications 9, 989 (2018).

42. Durisko, Z., Mulsant, B.H., McKenzie, K. & Andrews, P.W. Using evolutionary theory to guide mental health research. The Canadian Journal of Psychiatry 61, 159–165 (2016).

43. Mishra, G.D. et al. Early menarche, nulliparity and the risk for premature and early natural menopause. Human Reproduction 32, 679–686 (2017).

44. Ni, G. et al. Estimation of Genetic Correlation via Linkage Disequilibrium Score Regression and Genomic Restricted Maximum Likelihood. The American Journal of Human Genetics 102(2018).

